# TriGraphQA: a triple graph learning framework for model quality assessment of protein complexes

**DOI:** 10.64898/2026.03.17.712533

**Authors:** Luozhan Liang, Kailong Zhao

**Affiliations:** MOE Frontiers Science Center for Nonlinear Expectations, Research Center for Mathematics and Interdisciplinary Sciences, Shandong University, Qingdao 266237, China; Suzhou Research Institute of Shandong University, Suzhou, Jiangsu Province, 215123, China

**Author notes:** Corresponding authors: KZ.

**Keywords:** model quality assessment, protein complexes, graph neural network, triple-graph architecture

## Abstract

Accurate quality assessment of predicted protein-protein complex structures remains a major challenge. Existing graph-based quality assessment methods often treat the entire complex as a homogeneous graph, which obscures the physical distinction between intra-chain folding stability and inter-chain binding specificity. In this study, we introduce TriGraphQA, a novel triple graph learning framework designed for model quality assessment of protein complexes. TriGraphQA explicitly decouples monomeric and interfacial representations by constructing three geometric views: two residue-node graphs capturing the local folding environments of individual chains, and a dedicated contact-node graph representing the binding interface. Crucially, we propose an interface context aggregation module to project context-rich embeddings from the monomers onto the interface, effectively fusing multi-scale structural features. We conducted comprehensive tests on several challenging benchmark datasets, including Dimer50, DBM55-AF2, and HAF2. The results show that TriGraphQA significantly outperforms state-of-the-art single-model methods. TriGraphQA consistently achieves the highest global scoring correlations and lower top-ranking losses. Consequently, TriGraphQA provides a powerful evaluation tool for protein-protein docking, facilitating the reliable identification of near-native assemblies in large-scale structural modeling and molecular recognition studies.

## Introduction

Protein-protein interactions are the molecular engines driving numerous biological processes, ranging from signal transduction and metabolic pathways to complex immune responses[1–3]. Consequently, the accurate determination of protein complex structures is a cornerstone of modern structural biology and therapeutic design[4, 5]. With the advent of AlphaFold2 and the AlphaFold3, the field has recently undergone a paradigm shift, achieving unprecedented accuracy in predicting the structures of protein monomers and complexes[6–14]. Despite these significant advances, predicting the precise interfacial geometry of novel complexes, especially those involving antibody-antigen interactions or significant conformational changes, remains a formidable challenge[15, 16]. In these cases, computational docking is an indispensable tool for generating hypothetical binding postures[17–19]. However, docking procedures typically generate thousands of decoy models, creating a bottleneck known as the “scoring problem”: how to distinguish the key conformation from the large number of erroneous models[20, 21].

Traditionally, model quality assessment methods have primarily focused on individual proteins, relying on physics-based energy functions, stereochemistry checks, or machine learning scores, such as ModFOLD, DeepUMQA or ProQ series[22–31]. While these classic approaches have been specifically extended to address quaternary structures, capturing the complex physicochemical interactions at protein interfaces—such as specific residue stacking and oligomer stability—remains a challenging task[32]. To complement these single-model or quasi-single-model approaches, consensus-based methods are gaining increasing attention, especially when dealing with large numbers of docking decoys or predictive models[33–35]. For example, the MULTICOM series integrates multiple quality assessment scores, structural clustering, and pairwise model similarity to arrive at robust consensus rankings in selecting near-native conformations from diverse sets of multiple CASP experiments[36, 37]. However, consensus methods heavily depend on the diversity and quality of the input set, thus their performance degrades when dealing with large complexes, shallow or insufficiently informative multiple sequence alignments (e.g., in antibody-antigen systems), or situations where some excellent models exist but are ranked poorly[38].

In recent years, geometric deep learning, particularly graph neural networks (GNNs), has become the mainstream method for learning protein structure representations[39–41]. GNNs can naturally represent proteins as graphs, capturing the non-Euclidean geometry of residues and their interactions. Recently, several innovative methods have driven the development of this field. MViewEMA employs a multi-view learning framework, aggregating features from different structural representations to enhance the robustness of predictions[42]. ComplexQA leverages deep learning to integrate sequence and structural features, providing a comprehensive assessment of model quality[43]. TopoQA introduces topological features into the learning process, focusing on the persistence of structural gaps and loops to characterize complex stability[44]. Meanwhile, DProQA utilizes a specially designed gated graph converter architecture to rank candidate docking decoys by learning their relative quality[45]. Despite these advancements, existing approaches often treat the entire complex as a single homogeneous graph, which may obscure the distinction between intra-chain folding stability and inter-chain binding specificity. There is still significant room for improvement in how information is transferred from individual subunits to the interface. In this study, we propose a novel triple-graph neural network framework designed to explicitly decouple and then integrate intra-chain and inter-chain features for model quality assessment of protein complexes. Distinctively, our architecture characterizes the protein complex using three distinct geometric views: two residue-node graphs representing individual chains, and a dedicated contact-node graph representing the binding interface. Within the residue-node graphs, nodes correspond to residues and edges to residue contacts, capturing the local folding environment of each chain.

Conversely, in the contact-node graph, we adopt a dual representation where nodes denote inter-chain contacts and edges connect contacts sharing a common residue. Crucially, we employ a hierarchical pooling mechanism to aggregate context-rich embeddings from the residue-node graphs of the monomers and project them onto the contact-node graph of the interface. This allows the model to leverage the structural context of the participating monomers to assess interfacial interactions. By fusing these multi-scale features, our method effectively captures the synergy between monomer stability and interfacial complementarity, achieving superior performance in identifying high-quality docking models.

## Method

### 1. Overview of TriGraphQA

The overall workflow of TriGraphQA is illustrated in **Figure 1**. Given a 3D protein-protein complex model, TriGraphQA first extracts sequence and structural features to construct three decoupled geometric representations: two residue-node graphs for the individual chains and one contact-node graph for the binding interface. The monomeric graphs are independently processed through gated graph convolutional networks (GCNs) to capture the intra-chain folding environments and generate contextual embeddings. Concurrently, initial features of the contact-node graph are linearly transformed. Crucially, a dedicated interface context aggregation module projects the learned monomeric embeddings onto the corresponding interface contacts through a sequence-distance-aware weighting mechanism. The interface graph is subsequently processed by a final set of contact-node GCNs. A terminal linear layer then aggregates these multiscale representations to predict the overall model quality, outputting the final DockQ score.

**Figure 1.**
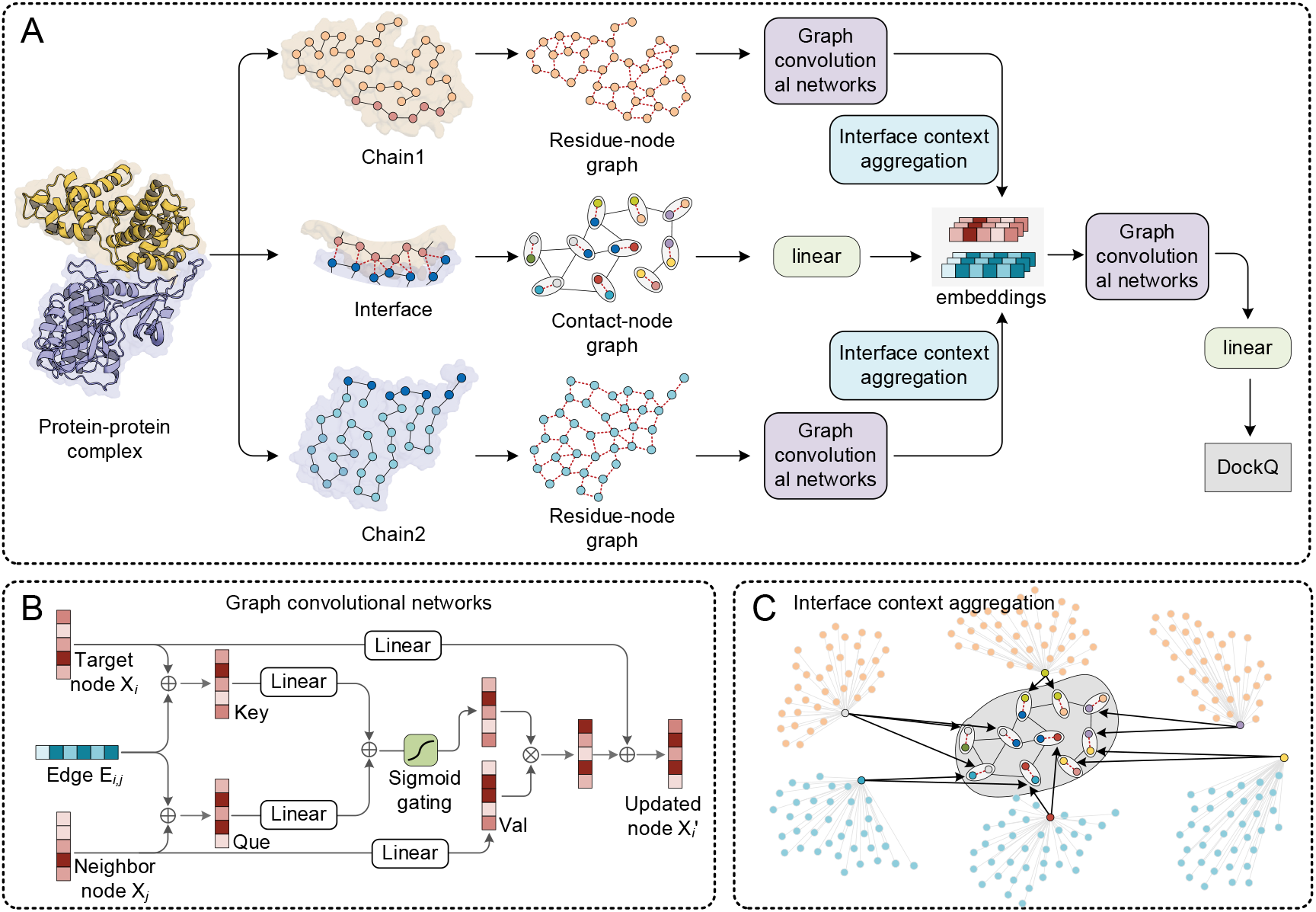
The framework of TriGraphQA. (A) Triple-graph architecture: two residue-node graphs for the individual chains and one contact-node graph for the binding interface, connected via the interface context aggregation module for synergistic feature fusion and DockQ prediction. (B) Gated graph convolution operation with edge-modulated message passing. (C) Interface context aggregation: projecting monomer folding contexts onto interface contacts through sequence-distance-aware weighting.

### 2. Dataset Construction

#### 2.1 Training datasets

To construct a robust and diverse dataset for training TriGraphQA, we established a comprehensive data pipeline consisting of data collection, decoy generation, and decoy sampling.

##### Data collection

We initially obtained non-redundant protein-protein complexes from the Q-BioLip database (with a cutoff date of January 2024)[46]. Then, we filtered the initial dataset, retaining only dimeric complexes with each monomer no longer than 500 amino acid residues, resulting in a dataset containing 10,502 unique protein dimers. *Decoy generation*. To enrich the structural space, we generated a diverse pool of decoy structures for each dimer using two approaches. First, we used AlphaFold3 to generate 25 decoy models for each target by varying the random seed[8]. Second, we used HDOCK to directly dock with the native structure, generating an additional 30 decoys[18].

##### Decoy sampling

To prevent model bias towards specific quality ranges and ensure balanced learning, we implemented a sampling strategy based on the true DockQ scores. The generated decoys were categorized into four quality bins: Incorrect (0 ≤ DockQ < 0.23), acceptable (0.23 ≤ DockQ < 0.49), medium quality (0.49 ≤ DockQ < 0.80), and high quality (0.80 ≤ DockQ ≤ 1.00)[47]. For each dimer, we sampled a maximum of 5 decoy structures from each bin, striving to keep the DockQ scores of the selected decoys as uniformly distributed as possible within each interval. This rigorous generation and sampling process ultimately produced a large-scale dataset containing 168,061 complex structures. From the dataset, we randomly selected 100 targets for the validation set and 50 targets for the benchmark set, denoted as Dimer50. To ensure the rigor of the evaluation and prevent data leakage, we removed all complexes with more than 40% sequence identity to the three test datasets (Dimer50, DBM55-AF2, and HAF2). After this redundancy filtering, the remaining 10,352 protein targets constituted the final training set.

#### 2.2 Test datasets

The test set includes three benchmark datasets: Dimer50, DBM55-AF2[45], and HAF2[43] dataset. For rigorous evaluation, we have ensured that the training set does not contain targets that are 40% sequence consistent with the test datasets.

### 3. Graph construction

To accurately capture the local folding environment of individual chains and the interaction details at the interface, TriGraphQA uses a novel triple graph architecture to represent a given protein-protein complex: two residue-node graphs for the individual chains and one contact-node graph dedicated to the binding interface.

#### 3.1 Residue-node graph for monomers

For each monomer, we construct a residue-node graph where nodes represent amino acid residues, with their 3D coordinates defined by the Cα atoms. If the minimum distance between any two heavy atoms of two residues is less than 5 Å, an edge is established between these two residues, representing a valid intrachain structural contact.

##### Node features (41 dimensions)

Each node is encoded with a 41-dimensional feature vector that integrates sequence and structural properties. Sequence-based features include a 20-dimensional one-hot encoding of the amino acid type and 7-dimensional Meiler features. Structure-based features consist of backbone bond lengths and angles (6 dimensions), Rosetta one-body energy terms (4 dimensions), and secondary structure classifications (4 dimensions), which include helix, sheet, coil, and an additional dimension specifically indicating missing or uncalculated assignments.

##### Edge features (21 dimensions)

Edges in the residue-node graph are characterized by a 21-dimensional vector. This includes five types of 2D distance maps (5 dimensions) adapted from DeepAccNet[48] and the spatial distance between the Cα atoms of the two residues (1 dimension), Rosetta two-body energy terms (7 dimensions), intra-chain hydrogen bond existence (2 dimensions), Euler rotation angles defining relative geometric orientations (6 dimensions), and the sequence distance (1 dimension), defined as the absolute index difference between the two connected residues along the primary sequence.

#### 3.2 Contact-node graph for the interface

Nodes in **c**ontact-node graph represent inter-chain residue contacts, defined as any pair of residues from different chains having at least one pair of heavy atoms within a distance of 7 Å. Edges are formed between two contact nodes if they share a common residue. Specifically, an edge connects the non-shared residues of two adjacent contacts.

##### Node features (37 dimensions)

Each contact node is parameterized by a 37-dimensional feature vector describing the properties of the interacting residue pair. This vector integrates four types of distances (4 dimensions) adapted from DeepAccNet (with the sequence separation feature excluded) and the spatial distance between the Cα atoms of the interacting residues (1 dimension), Euler rotation angles (6 dimensions), and the existence of intra- and inter-chain hydrogen bonds (4 dimensions). Furthermore, we introduce Rosetta two-body energy terms (7 dimensions), which are calculated for the entire complex and subsequently extracted for the specific interface residues. To capture the specific binding geometry, we incorporate an 8-dimensional encoding of the interacting residue pair and a 7-dimensional physical-aware relative orientation feature of the interacting residues, inspired by VoroIF-GNN[49, 50].

##### Edge features (26 dimensions)

The edges of the contact-node graph capture the higher-order geometric relationship between adjacent contacts. Following GraphPep[51], we calculate the spatial distance between the two non-shared residues and expand it into a 16-dimensional vector using radial basis functions (RBF) with centers evenly distributed from 0 to 15 Å and a width of 1.0. Additionally, the spatial angle formed by the triplet “non-shared residue 1 - shared residue center - non-shared residue 2” is calculated and discretized into 8 distinct bin sectors (8 dimensions). Finally, we added a two-dimensional positional feature to indicate the respective source chains of the interacting residues.

### 4. Network architecture

The overall architecture of TriGraphQA aims to process the structural information of protein-protein complexes through a parallel but interconnected triple graph framework. As illustrated in **Figure 1A**, the pipeline begins by independently feeding the residue-node graphs of chain 1 and chain 2 into their respective GCN modules. Simultaneously, the initial features of the contact-node graph are processed through a linear transformation. Following the monomeric feature extraction, the structural context of the individual chains is projected onto the interface via a novel interface context aggregation module. Finally, the fused context-rich embeddings are processed by another set of GCN modules dedicated to the contact-node graph, followed by a final linear layer to predict the overall quality of the docking model.

To effectively capture the complex interaction patterns within both the intra-chain and inter-chain environments, we use a gated graph convolutional network as the core building block of the graph. Inspired by the residual gated GCN framework[52], our message-passing mechanism explicitly incorporates edge features to modulate the information flow between neighboring nodes (**Figure 1B**).

Specifically, during the convolution process, the target node *X*_*i*_ and its neighbor node *X*_*j*_ are first fused with their connecting edge feature *E*_*i,j*_ through linear transformations to generate key and query, respectively. These representations are then summed and passed through a sigmoid activation function to produce an attention-like gating weight:

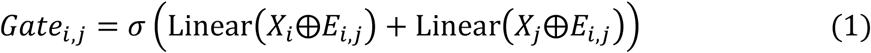

Where ⨁ denotes element-wise addition, and *σ* represents the sigmoid function. Simultaneously, the neighbor node *X*_*j*_ undergoes a linear transformation to generate a value vector. The gating weight is then used to perform an element-wise multiplication (⨂) with the value vector, filtering out spatial noise and retaining the most relevant structural features. Finally, the aggregated messages from all neighbors are combined with the original target node *X*_*i*_ via a residual skip connection to produce the updated node feature 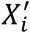. In our TriGraphQA framework, each GCN block in the monomer branches and the interface branch is composed of 3 consecutive layers of this gated convolution to ensure a sufficient receptive field.

### 5. Interface context aggregation module

To map the global folding context of the monomeric chains directly onto the local interfacial contacts, we propose the interface context aggregation module (**Figure 1C**).

For a given inter-chain contact node representing the interaction between residue 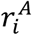 in monomer A and residue 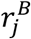 in monomer B, the prior 3-layer GCN modules transform the initial 41-dimensional node features of the residue-node graphs into 64-dimensional hidden representations. To encapsulate the structural support provided by monomer A to the interface residue 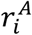, we aggregate the 64-dimensional features of all other residues in monomer A onto 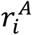. Crucially, the contribution of each residue 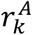 to the target residue 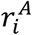 is determined by a carefully designed weighting function:

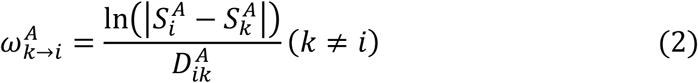

where 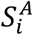 and 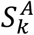 are the sequence indices of residues 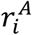 and 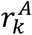, respectively, and 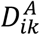 is the spatial Euclidean distance between their Ca atoms. Specifically, taking the natural logarithm of the sequence distance assigns higher importance to residues separated widely in the primary sequence, while the spatial distance denominator 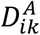 penalizes residues that are far apart in 3D space. Consequently, this weight selectively highlights long-range tertiary folding contacts that are spatially close to the interface residue, which are critical indicators of structural stability. The weighted features of all residues in monomer A are summed to form a comprehensive 64×1 context vector for residue 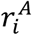. The identical procedure is independently applied to monomer B to generate a 64×1 context vector for residue 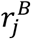.

Simultaneously, the initial 37-dimensional feature of the 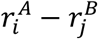 contact node is projected into a higher-dimensional space (128×1) using a linear layer. Finally, the aggregation module fuses these multiscale features by concatenating the linearly projected contact node feature with the context vectors of its constituent residues 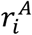 and 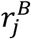:

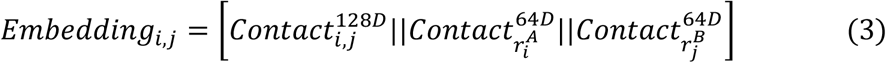

This operation yields a robust 256-dimensional (128±64±64) embedding for each contact node. By doing so, the updated contact-node graph is enriched with both high-resolution interfacial geometries and the global folding integrity of the monomers. These context-rich embeddings are subsequently fed into the final contact-node GCN layers to predict the complex quality.

## Results

### Performance Evaluation on the Dimer50 Dataset

To systematically evaluate the performance of TriGraphQA in protein-protein interaction quality assessment, we first benchmarked our model on the constructed Dimer50 dataset. As described in the Methods section, Dimer50 contains various 50 dimer complexes. For each target protein, we generated and extracted a model pool containing 55 decoys. On this dataset, we compared TriGraphQA with three complex model quality assessment methods: MViewEMA, TopoQA, and ComplexQA.

We first evaluated the global scoring accuracy of the methods by calculating the Pearson and Spearman correlation coefficients. It’s worth noting that TriGraphQA, TopoQA, and ComplexQA predict DockQ scores, while MViewEMA predict the global TM-score. Therefore, MViewEMA’s relevance metric is calculated between its predicted TM-score and the true TM-score, whereas the other methods use the DockQ score. Despite this difference in the target metrics, TriGraphQA demonstrated a substantial advantage over the other methods, achieving an average Pearson correlation of 0.534 and a Spearman correlation of 0.417. In contrast, MViewEMA (Pearson: 0.423, Spearman: 0.384), TopoQA (Pearson: 0.346, Spearman: 0.324), and ComplexQA (Pearson: 0.243, Spearman: 0.250) exhibited considerably lower correlation performance. The head-to-head scatter plots in **Figure 2A** further confirm this superiority.

**Figure 2.**
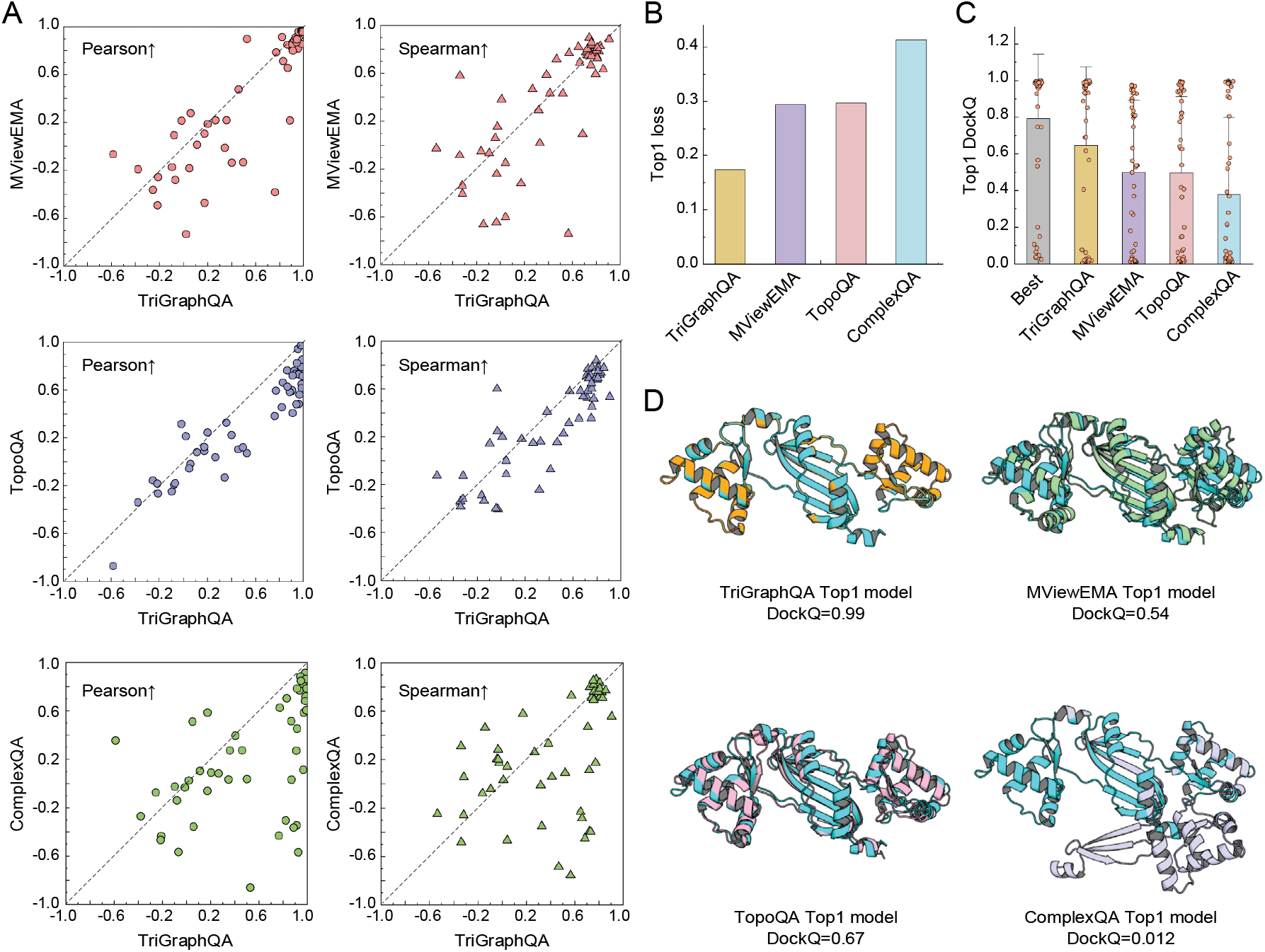
Performance comparison of TriGraphQA and other methods on the Dimer50 dataset. (A) Head-to-head comparison of the Pearson (circles) and Spearman (triangles) correlation coefficients for each target. Note that the correlations for TriGraphQA, TopoQA, and ComplexQA are calculated between the predicted and ground-truth DockQ scores, whereas the correlations for MViewEMA are calculated between the predicted and ground-truth global TM-score. (B) The average top-1 ranking loss of TriGraphQA and the three methods. (C) The distribution of the true DockQ scores for the top-1 models selected by each method. “Best” represents the theoretical upper bound, corresponding to the highest true DockQ score available in the decoy pool for each target. (D) Structural superpositions of the native structure and the top-1 models selected by different evaluation methods for the target 2FE3. The native structure is consistently colored in cyan.

In addition to overall correlation, the usefulness of a quality assessment tool largely depends on its ability to rank near-natural conformations at the top of the decoy pool. To evaluate this, we analyzed the top-1 ranking loss (**Figure 2B**) and the top-1 DockQ scores (**Figure 2C**). By ranking the decoys according to their respective predicted scores (TM-score for MViewEMA), TriGraphQA achieved the lowest top-1 ranking loss of 0.174, substantially outperforming MViewEMA (0.294), TopoQA (0.298), and ComplexQA (0.414). Consequently, the average DockQ score of the top-1 models selected by TriGraphQA reached 0.646. As illustrated in **Figure 2C**, where “Best” denotes the theoretical upper bound (the highest true DockQ score available in the decoy pool for each target), the distribution of TriGraphQA’s selected models is remarkably close to the optimal selections. Conversely, the other methods frequently fail to prioritize the best available near-native models, resulting in significantly lower average top-1 DockQ scores (0.499, 0.496, and 0.380 for MViewEMA, TopoQA, and ComplexQA, respectively).

To intuitively illustrate the effectiveness of TriGraphQA, **Figure 2D** presents a detailed case study on the target 2FE3. This target is the crystal structure of the Bacillus subtilis PerR-Zn[53], a metalloregulator that utilizes a novel Zn(Cys)4 structural redox switch to sense peroxide stress and regulate corresponding gene expressions. TriGraphQA successfully prioritized an outstanding near-native structure as its top-1 model, achieving a near-perfect DockQ score of 0.99. In contrast, MViewEMA and TopoQA selected models with noticeable structural and interfacial deviations (DockQ = 0.54 and 0.67, respectively), while ComplexQA completely failed to recognize the correct binding pattern, selecting an incorrect decoy with a DockQ score of 0.012. This specific example highlights TriGraphQA’s exceptional capacity to capture key interfacial geometries and accurately score complex biological interactions.

### Performance Evaluation on the DBM55-AF2 Dataset

To further evaluate the robustness and generalizability of TriGraphQA, we extended our evaluation to the DBM55-AF2 benchmark[45]. This dataset, constructed by Jianlin Cheng’s team in their DProQA research, consists of multimeric decoys generated by AlphaFold-Multimer. It provides a rigorous testbed for evaluating scoring functions on predicted structures. Since TriGraphQA is inherently designed for dimeric interactions, we adapted our approach for multimers by decomposing each complex into its constituent pairwise dimers. Specifically, the overall predicted DockQ score for a multimeric decoy is calculated by averaging the predicted scores of its valid interacting pairs, utilizing the native structure to identify which pairs are truly interacting.

The global correlation results are shown in **Figure 3A**. It is worth noting that using native topological information to filter and average effective interaction pairs may have a slight advantage in global correlation analysis (Pearson and Spearman) compared to other methods that evaluate the entire polymer without such prior knowledge. Nevertheless, despite this disparity in scoring protocols, TriGraphQA maintains its robust global scoring capability on this challenging dataset. It achieves an average Pearson correlation of 0.421, consistently outperforming TopoQA (0.309), DProQA (0.195), ComplexQA (−0.127), and MViewEMA (0.369). In terms of Spearman correlation (0.289), our method matches MViewEMA while substantially surpassing the remaining methods. The scatter plots confirm this target-level superiority, with the majority of individual data points falling below the reference diagonal.

**Figure 3.**
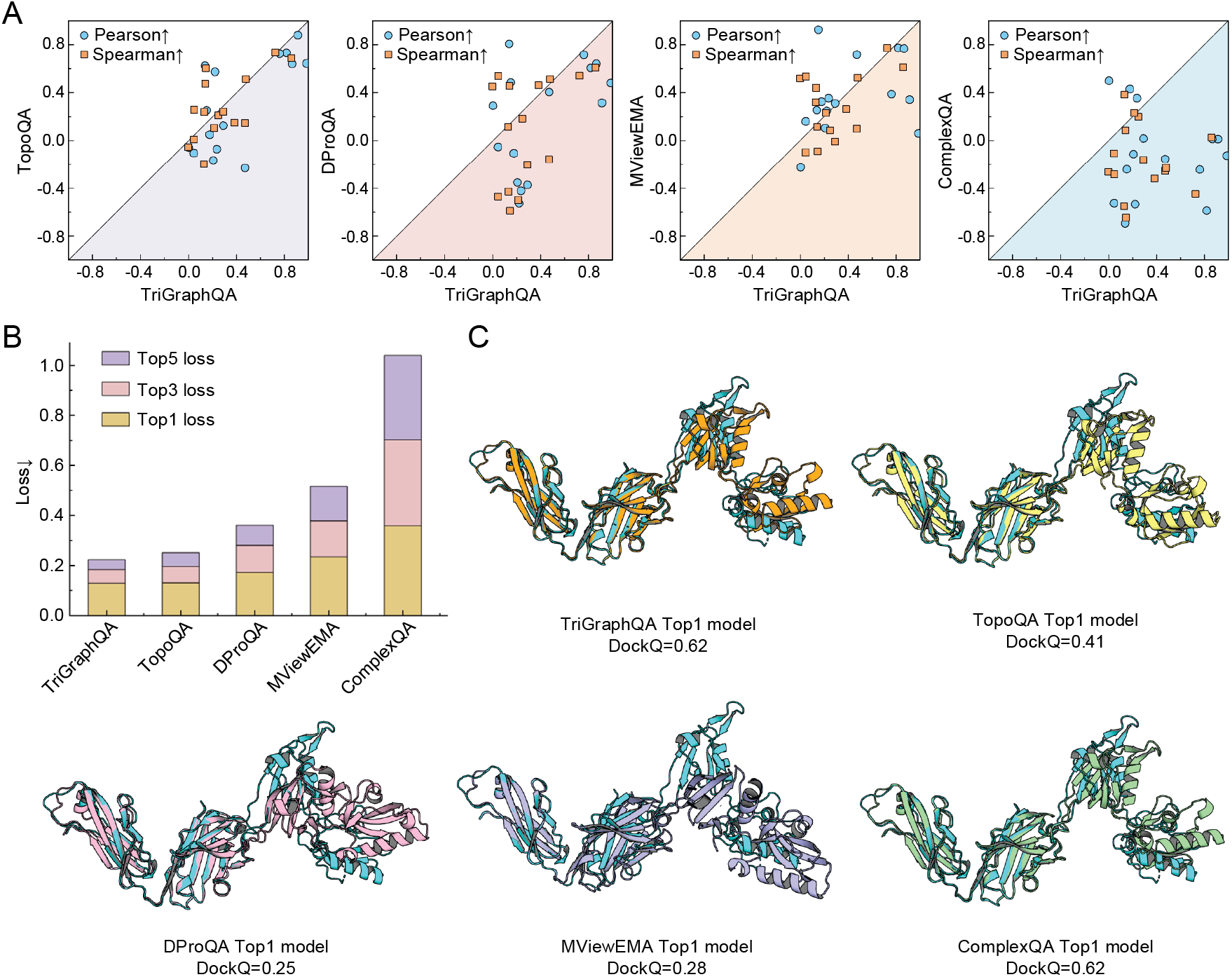
Performance comparison of TriGraphQA and other methods on the DBM55-AF2 dataset. **(A)** Head-to-head comparison of the Pearson and Spearman correlation between the predicted and ground-truth DockQ scores for each target. Points located below the diagonal indicate targets where TriGraphQA achieves a higher correlation score than the corresponding method. **(B)** The average top-1, top-3, and top-5 ranking loss of TriGraphQA and the four baseline methods across the dataset, represented as stacked bars. A lower ranking loss indicates a better capability to retrieve near-native models from the decoy pool. **(C)** Structural superpositions of the native structure and the top-1 models selected by different evaluation methods for the target 6AL0. The native structure is consistently colored in blue (cyan).

**Figure 4.**
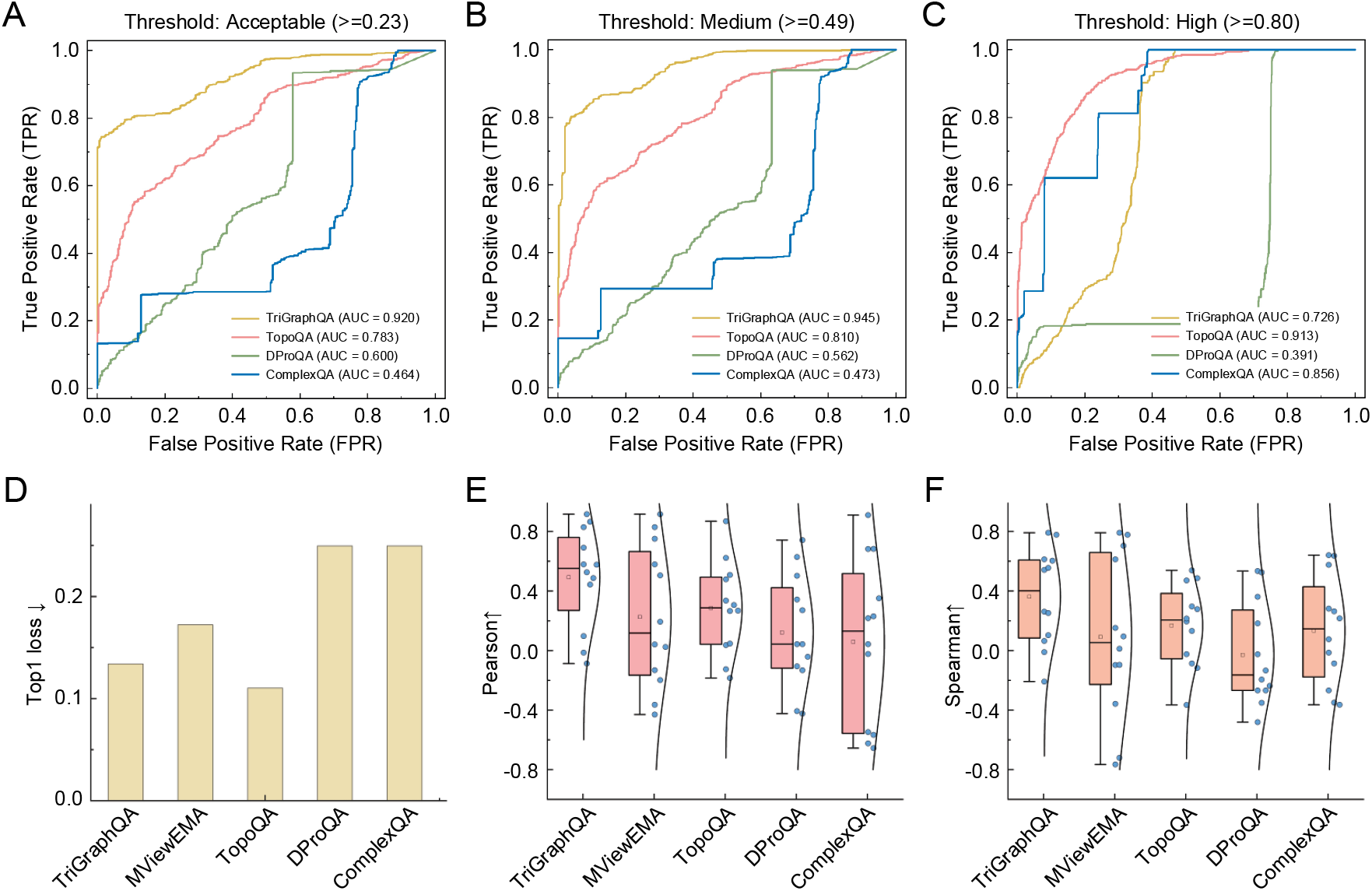
Performance evaluation of TriGraphQA and other methods on the HAF2 dataset. **(A-C)** Receiver operating characteristic (ROC) curves evaluating the ability of the methods to distinguish decoys at three DockQ quality cutoffs. MViewEMA is excluded from these plots as it predicts global TM-scores. **(D)** Bar chart comparing the average top-1 ranking loss of TriGraphQA against MViewEMA, TopoQA, DProQA, and ComplexQA. **(E-F)** The distribution of Pearson and Spearman correlation coefficients for each target. Each blue dot represents a target.

Crucially, while our calculation of the total score for multimers relies on native pairing information, it acts as a constant linear transformation for all decoys of a given target and thus does not bias the relative ranking within the decoy pool. In terms of top-level model selection performance, TriGraphQA successfully minimizes the ranking losses across multiple thresholds (**Figure 3B**). It records an average top-1 loss of 0.128, a top-3 loss of 0.055, and a top-5 loss of 0.039, strictly outperforming all four competitive methods. Consequently, the average DockQ score of the top-1 models identified by our method stands at 0.487, maintaining a steady lead over the top selections made by TopoQA (0.474), DProQA (0.432), MViewEMA (0.369), and ComplexQA (0.246).

A structural visualization of target 6AL0 (**Figure 3C**) further demonstrates the accuracy of our model. This target represents an antibody-antigen complex (the NZ-1 Fab fragment bound to a PDZ tandem fragment), which is notoriously difficult to score due to the high flexibility and specificity required at the binding interface. Despite these difficulties, TriGraphQA successfully retrieved a high-quality near-native structure (DockQ = 0.62) as its top rank. While ComplexQA also managed to pick a model with same score for this specific target, the selections made by TopoQA, MViewEMA, and DProQA exhibited pronounced interfacial deviations. This further confirms the advantage of TriGraphQA’s triple graph structure in capturing interfacial complementarity, even in highly challenging biological systems.

### Performance Evaluation on the HAF2 Dataset

To further evaluate the model’s applicability to heterodimeric complexes, we evaluated TriGraphQA using the HAF2 dataset[43]. This dataset, created by Renzhi Cao’s team for ComplexQA research, was initially generated by AlphaFold-Multimer and contains 1370 decoy models distributed across 13 heterodimeric targets. To ensure fairness in the comparison and avoid overestimating model performance, we followed TopoQA and selected 12 valid heterodimeric targets for subsequent evaluation.

We first evaluated the model’s ability to distinguish between valid binding poses and false decoys at different DockQ thresholds. Receiver operating characteristic (ROC) curves were plotted to distinguish decoys at three standard DockQ quality thresholds: acceptable (⩾0.23), medium (⩾0.49), and high (⩾0.80) (**Figures 3A-C**). Note that MViewEMA was excluded from this specific analysis as it predicts global TM-score rather than DockQ scores. TriGraphQA demonstrates exceptional assessment performance, achieving the highest Area under curve (AUC) for both the acceptable (AUC=0.920) and medium (AUC=0.945) quality thresholds, significantly outperforming TopoQA, DProQA, and ComplexQA. While TopoQA and ComplexQA show strong AUCs at the extremely strict High-quality threshold (**Figure 3C**), TriGraphQA’s superior performance across the broader acceptable and medium ranges indicates a more robust ability to filter out fundamentally incorrect docking models from vast decoy pools.

Finally, we examined the absolute ranking errors and the global scoring correlations (**Figures 3D-F**). Although TopoQA achieves a marginally lower average top-1 ranking loss (0.110) compared to TriGraphQA (0.134), our method demonstrates a significant superiority in maintaining target-level scoring consistency. As depicted in the violin plots for Pearson and Spearman correlations, TriGraphQA not only achieves the highest average correlation coefficients (Pearson: 0.493, Spearman: 0.362) but also exhibits a much more concentrated and stable distribution. In contrast, the other methods display highly dispersed distributions with numerous targets falling into negative correlation territories. Furthermore, we analyzed the top-10 hit rate, defined as the total number of qualified decoys that successfully ranked in the top-10 across all targets. While all methods performed comparablely in retrieving the most stringent high-quality models, TriGraphQA successfully evaluated and retrieved the most decoys at both medium and acceptable thresholds. This underscores the advantage of TriGraphQA’s multi-scale graph representation in consistently estimating the relative quality of heterogeneous docking models.

### Ablation Study

To investigate the essential contribution of our proposed triple-graph architecture and the interface context aggregation module, we conducted an ablation study on the Dimer50 dataset, comparing the TriGraphQA model with two architectural variants: TriGraphQA-1 and TriGraphQA-2. Specifically, TriGraphQA-1 employs a dual-graph architecture. Instead of explicitly decoupling the monomers, it constructs a global graph representing the entire complex and a local graph representing only the interface. These two graphs are processed independently through GCN to predict separate DockQ scores, which are then combined via a weighted average to produce the final score. On the other hand, TriGraphQA-2 utilizes a single-graph architecture that exclusively models the interface graph, completely discarding the global structural context of the individual chains.

As summarized in **Table 1**, the TriGraphQA model substantially outperforms both variants across all evaluation metrics. When examining TriGraphQA-2, we observe a significant decrease in its ability to prioritize optimal conformations, resulting in a top-1 ranking loss of up to 0.335. Its Pearson correlation (0.482) and Spearman correlation (0.315) remain relatively competitive, which indicates that interface-specific features are indeed highly informative for general quality assessment. Nevertheless, the poor ranking loss reveals that neglecting the intra-chain folding stability of individual monomers severely compromises the model’s precision in identifying the absolute near-native conformation among top candidates.

**Table 1.** The results of ablation study.

|  | Top1 loss | Pearson | Spearman |
| --- | --- | --- | --- |
| TriGraphQA | 0.173 | 0.534 | 0.417 |
| TriGraphQA-1 | 0.288 | 0.410 | 0.262 |
| TriGraphQA-2 | 0.335 | 0.482 | 0.315 |

More interestingly, the performance of TriGraphQA-1 demonstrates that simply integrating global contextual information is insufficient. By incorporating the entire complex into a single global graph and performing late-stage score averaging, TriGraphQA-1’s overall correlation significantly decreases, and its top-1 loss is much higher than that of the complete model. This result indicates that treating the entire complex as a homogeneous graph or fusing global and local information only at the scalar score stage is inadequate. By explicitly separating interacting monomers from interface regions and fusing their contextual embeddings at the feature level, the relationship between monomer folding stability and interface complementarity can be captured more effectively, resulting in optimal overall prediction performance.

## Conclusion

This study proposes a novel multi-scale geometric deep learning framework, TriGraphQA, based on a triple graph architecture for model quality assessment of protein-protein complexes. Recognizing the limitations of existing approaches that treat protein complexes as homogeneous graphs, TriGraphQA explicitly decouples intra-chain folding stability from inter-chain binding specificity. To achieve this, we constructed a unique representation that comprises two residue-node graphs for individual monomers and a dedicated contact-node graph for the binding interface. Combined with a specially designed interface context aggregation module, our method effectively projects the global structural context of monomeric chains onto interfacial geometries.

We conducted comprehensive evaluations on several challenging benchmark datasets, including Dimer50, DBM55-AF2, and HAF2. The results clearly demonstrate that TriGraphQA significantly outperforms the state-of-the-art methods such as MViewEMA, TopoQA, and ComplexQA. TriGraphQA consistently achieved the highest global scoring correlations (Pearson and Spearman) and substantially lower ranking errors for top models. Notably, even in difficult cases such as multi-chain assemblies and highly flexible protein complexes, our method exhibited strong robustness and high accuracy in reliably selecting near-native conformations from large decoy pools. Furthermore, ROC curve analysis further confirmed its excellent performance in filtering out clearly incorrect docking poses across different quality thresholds.

In summary, by effectively capturing the multi-scale relationship between individual subunits and their binding interfaces, TriGraphQA offers a highly accurate and reliable tool for assessing the quality of protein complex models. While the current results are promising, TriGraphQA handles multimeric assemblies by breaking them down into pairwise dimers, which may not fully capture the overall, higher-order interactions in very large macromolecular complexes. Future improvements will focus on extending this framework to directly model true multi-body interactions and optimizing its computational efficiency for ultra-large protein complexes. We hope this approach will ultimately help advance computational docking pipelines and support more accurate structural predictions in drug design and complex biological studies.

## Data availability

The benchmark dataset can be downloaded at https://huggingface.co/datasets/Luozhan/Dimer50/tree/main

## Software availability

The source code and model parameters are available at https://github.com/1874427280/TriGraphQA

## Acknowledgements

We thank Jianyi Yang for his discussions and feedback. This work is supported by the following funding sources: National Natural Science Foundation of China (NSFC 62503282), the Basic Research Program of Jiangsu Province (BK20250432), Postdoctoral Fellowship Program and China Postdoctoral Science Foundation (2025M773121), Qingdao Postdoctoral Applied Research Project, and Fundamental Research Funds for the Central Universities.

## Reference

1. Rao VS, Srinivas K, Sujini G et al. Protein-protein interaction detection: methods and analysis, International journal of proteomics 2014;2014:147648.

2. Jones S, Thornton JM. Principles of protein-protein interactions, Proceedings of the National Academy of Sciences 1996;93:13–20.

3. Gao Z, Jiang C, Zhang J et al. Hierarchical graph learning for protein–protein interaction, Nature communications 2023;14:1093.

4. Greenblatt JF, Alberts BM, Krogan NJ. Discovery and significance of protein-protein interactions in health and disease, Cell 2024;187:6501–6517.

5. Ebrahimi SB, Samanta D. Engineering protein-based therapeutics through structural and chemical design, Nature communications 2023;14:2411.

6. Jumper J, Evans R, Pritzel A et al. Highly accurate protein structure prediction with AlphaFold, Nature 2021;596:583–589.

7. Baek M, DiMaio F, Anishchenko I et al. Accurate prediction of protein structures and interactions using a three-track neural network, Science 2021;373:871–876.

8. Abramson J, Adler J, Dunger J et al. Accurate structure prediction of biomolecular interactions with AlphaFold 3, Nature 2024;630:493–500.

9. Evans R, O’neill M, Pritzel A et al. Protein complex prediction with AlphaFold-Multimer, biorxiv 2021:2021.2010. 2004.463034.

10. Guo Z, Liu J, Skolnick J et al. Prediction of inter-chain distance maps of protein complexes with 2D attention-based deep neural networks, Nature communications 2022;13:6963.

11. Wang W, Luo Y, Peng Z et al. Accurate Biomolecular Structure Prediction in CASP16 With Optimized Inputs to State - Of - The - Art Predictors, Proteins: Structure, Function, and Bioinformatics 2026;94:142–153.

12. Peng Z, Wang W, Wei H et al. Improved protein structure prediction with trRosettaX2, AlphaFold2, and optimized MSAs in CASP15, Proteins: Structure, Function, and Bioinformatics 2023;91:1704–1711.

13. Zhao K, Xia Y, Zhang F et al. Protein structure and folding pathway prediction based on remote homologs recognition using PAthreader, Communications biology 2023;6:243.

14. Zhao K, Zhao P, Wang S et al. FoldPAthreader: predicting protein folding pathway using a novel folding force field model derived from known protein universe, Genome Biology 2024;25:152.

15. Yin R, Pierce BG. Evaluation of AlphaFold antibody–antigen modeling with implications for improving predictive accuracy, Protein Science 2024;33:e4865.

16. Fernández-Quintero ML, Kokot J, Waibl F et al. Challenges in antibody structure prediction. In: MAbs. 2023, p. 2175319. Taylor & Francis.

17. Vakser IA. Protein-protein docking: From interaction to interactome, Biophysical journal 2014;107:1785–1793.

18. Yan Y, Tao H, He J et al. The HDOCK server for integrated protein–protein docking, Nature protocols 2020;15:1829–1852.

19. Honorato RV, Trellet ME, Jiménez-García B et al. The HADDOCK2. 4 web server for integrative modeling of biomolecular complexes, Nature protocols 2024;19:3219–3241.

20. Halperin I, Ma B, Wolfson H et al. Principles of docking: An overview of search algorithms and a guide to scoring functions, Proteins: Structure, Function, and Bioinformatics 2002;47:409–443.

21. Liang F, Sun M, Xie L et al. Recent advances and challenges in protein complex model accuracy estimation, Computational and Structural Biotechnology Journal 2024;23:1824–1832.

22. Guo S-S, Liu J, Zhou X-G et al. DeepUMQA: ultrafast shape recognition-based protein model quality assessment using deep learning, Bioinformatics 2022;38:1895–1903.

23. McGuffin LJ. The ModFOLD server for the quality assessment of protein structural models, Bioinformatics 2008;24:586–587.

24. Uziela K, Menéndez Hurtado D, Shu N et al. ProQ3D: improved model quality assessments using deep learning, Bioinformatics 2017;33:1578–1580.

25. Adhikari S, Schop M, de Boer IJ et al. Protein quality in perspective: a review of protein quality metrics and their applications, Nutrients 2022;14:947.

26. Zhang L, Ding R, Chen X et al. ContrastQA: A label-guided graph contrastive learning-based approach for protein complex structure quality assessment, biorxiv 2025:2025.2006. 2020.660832.

27. Baldassarre F, Menéndez Hurtado D, Elofsson A et al. GraphQA: protein model quality assessment using graph convolutional networks, Bioinformatics 2021;37:360–366.

28. Cao R, Bhattacharya D, Hou J et al. DeepQA: improving the estimation of single protein model quality with deep belief networks, BMC bioinformatics 2016;17:495.

29. Olechnovič K, Venclovas Č. VoroMQA: Assessment of protein structure quality using interatomic contact areas, Proteins: Structure, Function, and Bioinformatics 2017;85:1131–1145.

30. Roney JP, Ovchinnikov S. State-of-the-art estimation of protein model accuracy using AlphaFold, Physical review letters 2022;129:238101.

31. Guo L, He J, Lin P et al. TRScore: a 3D RepVGG-based scoring method for ranking protein docking models, Bioinformatics 2022;38:2444–2451.

32. Fadini A, Studer G, Read RJ. Model quality assessment for CASP16, Proteins: Structure, Function, and Bioinformatics 2026;94:302–313.

33. Roy RS, Liu J, Giri N et al. Combining pairwise structural similarity and deep learning interface contact prediction to estimate protein complex model accuracy in CASP15, Proteins: Structure, Function, and Bioinformatics 2023;91:1889–1902.

34. McGuffin LJ, Alhaddad SN, Behzadi B et al. Prediction and quality assessment of protein quaternary structure models using the MultiFOLD2 and ModFOLDdock2 servers<? mode pagerangestyle?>, Nucleic Acids Research 2025;53:W472–W477.

35. Liu D, Liu J, Wang H et al. DeepUMQA-X: Comprehensive and insightful estimation of model accuracy for protein single-chain and complex, Nucleic Acids Research 2025;53:W219–W227.

36. Liu J, Guo Z, Wu T et al. Improving AlphaFold2-based protein tertiary structure prediction with MULTICOM in CASP15, Communications chemistry 2023;6:188.

37. Liu J, Wu T, Guo Z et al. Improving protein tertiary structure prediction by deep learning and distance prediction in CASP14, Proteins: Structure, Function, and Bioinformatics 2022;90:58–72.

38. Shuvo MH, Bhattacharya S, Bhattacharya D. QDeep: distance-based protein model quality estimation by residue-level ensemble error classifications using stacked deep residual neural networks, Bioinformatics 2020;36:i285–i291.

39. Fout A, Byrd J, Shariat B et al. Protein interface prediction using graph convolutional networks, Advances in neural information processing systems 2017;30.

40. Bronstein MM, Bruna J, LeCun Y et al. Geometric deep learning: going beyond euclidean data, IEEE Signal Processing Magazine 2017;34:18–42.

41. Réau M, Renaud N, Xue LC et al. DeepRank-GNN: a graph neural network framework to learn patterns in protein–protein interfaces, Bioinformatics 2023;39:btac759.

42. Liu D, Zhao X, Zhang T et al. MViewEMA: Efficient Global Accuracy Estimation for Protein Complex Structural Models Using Multi-View Representation Learning, biorxiv 2025:2025.2007. 2025.666906.

43. Zhang L, Wang S, Hou J et al. ComplexQA: a deep graph learning approach for protein complex structure assessment, Briefings in bioinformatics 2023;24:bbad287.

44. Han B, Zhang Y, Li L et al. TopoQA: a topological deep learning-based approach for protein complex structure interface quality assessment, Briefings in bioinformatics 2025;26:bbaf083.

45. Chen X, Morehead A, Liu J et al. A gated graph transformer for protein complex structure quality assessment and its performance in CASP15, Bioinformatics 2023;39:i308–i317.

46. Wei H, Wang W, Peng Z et al. Q-biolip: A comprehensive resource for quaternary structure-based protein–ligand interactions, Genomics, Proteomics and Bioinformatics 2024;22:qzae001.

47. Basu S, Wallner B. DockQ: a quality measure for protein-protein docking models, PloS one 2016;11:e0161879.

48. Hiranuma N, Park H, Baek M et al. Improved protein structure refinement guided by deep learning based accuracy estimation, Nature communications 2021;12:1340.

49. Olechnovič K, Venclovas Č. VoroIF-GNN: Voronoi tessellation-derived protein–protein interface assessment using a graph neural network, Proteins: Structure, Function, and Bioinformatics 2023;91:1879–1888.

50. Xie J, Zhang Y, Wang Z et al. PPI-Graphomer: enhanced protein-protein affinity prediction using pretrained and graph transformer models, BMC bioinformatics 2025;26:116.

51. Tao H, Wang X, Huang S-Y. An interaction-derived graph learning framework for scoring protein– peptide complexes, Nature Machine Intelligence 2025:1–12.

52. Bresson X, Laurent T. Residual gated graph convnets, arXiv preprint 1711.07553 2017.

53. Traoré DA, El Ghazouani A, Ilango S et al. Crystal structure of the apo-PerR-Zn protein from Bacillus subtilis, Molecular microbiology 2006;61:1211–1219.

